# Common fluorescent *Pseudomonas* in the phyllosphere can influence aphid behavior in diverse ways

**DOI:** 10.1101/2024.04.26.591271

**Authors:** Kathryn L Herr, Jonah Schieber, Tory A Hendry

**Author notes:** Corresponding Author: Tory A Hendry.

## Abstract

Bacteria in the phyllosphere, the above ground parts of plants, can have complex interactions with plants and their insect visitors. For instance, certain *Pseudomonas* strains change pea aphid (*Acyrthosiphon pisum*) feeding behavior. Aphids visually detect and avoid feeding when *Pseudomonas* produces a compound with blue fluorescent emissions, the siderophore pyoverdine. It is unknown how commonly such interactions occur in nature, so we investigated this potential in pea plant (*Pisum sativum*) phyllosphere communities. We found a diversity of *Pseudomonas* taxa with fluorescent potential in pea plant microbiomes. Culture isolates revealed a wide range of fluorescent emissions spectra across taxa, which produced blue to green fluorescence of varying intensities. We tested strains from across this emissions range for pea aphid behavioral responses when given a choice between plants inoculated with a fluorescent isolate or a control treatment. Consistent with previous work, we found that some isolates were avoided by aphids. Surprisingly, some isolates were actually attractive to aphids, causing up to 70% of aphids to settle and feed on bacterially treated plants. Attractive isolates produced green fluorescence in culture, suggesting that attraction could be due to aphid sensory biases. Overall, we found both isolate fluorescence in culture and siderophore potential inferred from gene content to be poor predictors of aphid behavior. We also found very high variability in responses across replicate experiments for some strains, suggesting that environmental conditions may influence outcomes. This shows that bacterial fluorescence may be common on plants and can have context-dependent impacts on herbivorous insects.

## Introduction

The phyllosphere, or the above ground parts of plants, is known to house diverse interacting organisms, including microbes and herbivorous insects (Humphrey et al. 2014; Bashir et al. 2022). Bacteria in the genus *Pseudomonas* (pseudomonads) are common members of phyllosphere communities and are often studied for their direct impacts on plants as pathogens or beneficial taxa (Passera et al. 2019; Abdellatif et al. 2022; Agbavor et al. 2022). However, interactions of pseudomonads with herbivorous insects can also have important ecological outcomes (Humphrey et al. 2014; Smee and Hendry 2022). For instance, plant-associated *Pseudomonas* can influence both the survival and behavior of aphid pest insects (Stavrinides et al. 2009; Hendry et al. 2016; Smee et al. 2021; Smee and Hendry 2022; Silva-Sanzana et al. 2023). Aphids avoid feeding on leaves with epiphytic populations of certain *Pseudomonas syringae* strains due to fluorescent emissions from bacterially produced compounds (Hendry et al. 2018). Fluorescent pseudomonads in the phyllosphere therefore have the potential to influence aphid distributions and feeding choices in nature. However, it is unknown how prevalent fluorescent compound production is in natural phyllosphere microbial communities. Fluorescent pseudomonad interactions that influence aphid pest populations and behavior are especially relevant to plant health and agricultural sustainability, yet are poorly understood.

Pseudomonads are genetically and phenotypically diverse and inhabit a wide range of environments, including soil and host-associated habitats (Morris et al. 2013; Saati-Santamaría et al. 2022). They include commensals and pathogens of both plants and animals, and several clades of pseudomonads are commonly plant-associated (Rai et al. 2017; Passera et al. 2019; Shalev, Karasov, et al. 2022; Shalev, Ashkenazy, et al. 2022). These are predominantly taxa within the species complexes *P. syringae* and *P. fluorescens* (Choudhary et al. 2009; Passera et al. 2019). Some of these phyllosphere dwelling taxa are considered plant beneficial because they promote plant growth or protect against pathogens (Byrne et al. 2005; Ye, Hildebrand, et al. 2014; Zengerer et al. 2018; Legein et al. 2020; Santos Kron et al. 2020; Shalev, Ashkenazy, et al. 2022; Mehlferber et al. 2023).These bacteria can also protect against insects (Humphrey et al. 2014; Flury et al. 2016). For example, strains in the species complex *P. syringae*, as well as some other *Pseudomonas* taxa, can be highly infective and virulent to aphids (Hendry et al. 2016; Smee et al. 2017; Smee et al. 2021), and therefore have promise as biological control agents (Gerhardson 2002; Paliwal et al. 2022). Aphids are sap sucking insects and major vectors of serious plant pathogens (Ng and Perry 2004). They can cause damage and crop loss in agriculture but can be difficult to control (El Fakhouri et al. 2021; Dhillon et al. 2022; Luo et al. 2022; Ali et al. 2023). Plant beneficial bacteria present an opportunity for novel and sustainable aphid control strategies, such as using fluorescent pseudomonads to deter aphids (Hendry et al. 2018). However, we currently lack an understanding of how commonly fluorescent pseudomonads interact with aphids on plants, as well as the range of outcomes from these interactions.

Pseudomonads often naturally produce green or blue fluorescent emissions, with fluorescence primarily caused by iron scavenging compounds called siderophores (Ferret et al. 2014; Schalk et al. 2020), or other compounds such as phenazine (Chin-A-Woeng et al. 1998; Wang et al. 2011; Mavrodi et al. 2013; Vilaplana and Marco 2020). When excited by ultraviolet (UV) light, the siderophore pyoverdine emits visible light (Meyer and Abdallah 1978; Mureseanu et al. 2003; del Olmo et al. 2003). These light emissions from *Pseudomonas* can alter the spectra of the leaf in ways that are visually detectable by aphids (Döring and Chittka 2007; Hendry et al. 2018). Siderophores are a functional class of compounds, and a diversity of types are typically produced by pseudomonads (Ahmed and Holmström 2014; Schalk et al. 2020). Some of these, such as pyoverdine and pyochelin, are fluorescent due to an iron binding chromophore that is absent in non-fluorescent siderophores (Bultreys et al. 2001). Pyoverdine, the most common fluorescent siderophore produced by *Pseudomonas* (Budzikiewicz 1997; Meyer 2000), can vary in emission spectra from blue green to yellow green (wavelengths from 400 – 550 nm) (Elliott 1958; Meyer 2000; Tank et al. 2012). In addition to being influenced by siderophore structure, the fluorescent emission color of siderophores is also influenced by whether they are bound to iron (Meyer and Abdallah 1978; Greenwald et al. 2008; Hendry et al. 2018) and the pH of the environment (Xiao and Kisaalita 1995).

Siderophores are excreted by microbes when iron is limiting and bind environmental iron, sequestering it for subsequent cellular reuptake. Production of siderophores may be favorable on plant surfaces due to low iron availability (Legein et al. 2020), but epiphytic bacteria on plant surfaces don’t produce these compounds under all conditions, and which conditions drive siderophore production in the phyllosphere is poorly understood (Joyner and Lindow 2000; Karamanoli et al. 2011; Yu et al. 2013; Helmann et al. 2019). In addition to iron limitation, availability of various carbon sources and also temperature may influence siderophore production in plant-associated pseudomonads (Garibaldi 1971; Duffy and Défago 1999; Owen and Ackerley 2011; Arvizu-Gómez et al. 2013; Mendonca et al. 2020; Tribelli and López 2022). Additionally, while pyoverdine is the primary siderophore of fluorescent P*seudomonas*, pseudomonads are often capable of producing secondary siderophores such as nonfluorescent achromobactin (Cornelis and Matthijs 2002; Jones et al. 2007; Owen and Ackerley 2011; Schalk et al. 2020). Shifts to production of nonfluorescent siderophores over pyoverdine would alter fluorescent emissions of bacteria in the phyllosphere. The production of other fluorescent compounds like phenazine is also influenced by environmental factors (Serafim et al. 2023), making fluorescent emissions by bacteria on plants difficult to predict.

Previous work found that pea aphids (*Acyrthosiphon pisum*) avoided epiphytic bacterial strains of *P. syringae* that shifted the color of leaves to be more blue with fluorescent emissions (Hendry et al. 2018). This blue fluorescence was due to production of pyoverdine, and strains that could not produce pyoverdine, or that did not change leaf color as much, were not avoided (Hendry et al. 2018). This study did not include other colors of bacterial fluorescence or a diversity of pseudomonads. Pseudomonads are common in the phyllosphere, and diversity in the color and frequency of their fluorescent emissions likely influences whether any specific strain or leaf microbial community alters aphid behavior.

Here, we use sequence independent and dependent approaches to investigate the diversity and abundance of pseudomonads in the snap pea (*Pisum sativum*) phyllosphere, as well as the range of fluorescence colors these strains produce. We metagenomically sequenced pea phyllosphere communities and genomically sequenced pseudomonad isolates to define the fluorescence potential of these communities. Peas are an important crop plant but the phyllosphere microbiome of these plants has not been investigated. Peas regularly interact with pea aphids, which we use here as a model species in behavioral experiments with pseudomonad isolates. To confirm that blue fluorescence from a diversity of pseudomonads deters aphids, and to test for behavioral outcomes with other fluorescence colors, we conducted choice experiments using isolates that vary in their fluorescence spectra. This study is the first to explore the broad potential of epiphytic *Pseudomonas* to influence aphid behavior in the phyllosphere.

## Methods

### Sample collection and processing

To investigate the epiphytic bacterial community of *P. sativum* leaves, we collected snap pea leaves at three agricultural sites in Ithaca, NY during the summer of 2019 (Table 1). Leaves were sampled from Here We Are Farm (HWAF), Dilmun Hill Student Farm (DHSF), and Stick and Stone Farm (SASF). Humidity and temperature were recorded at each collection site using a Thermpro Digital Thermometer.

**Table 1.**
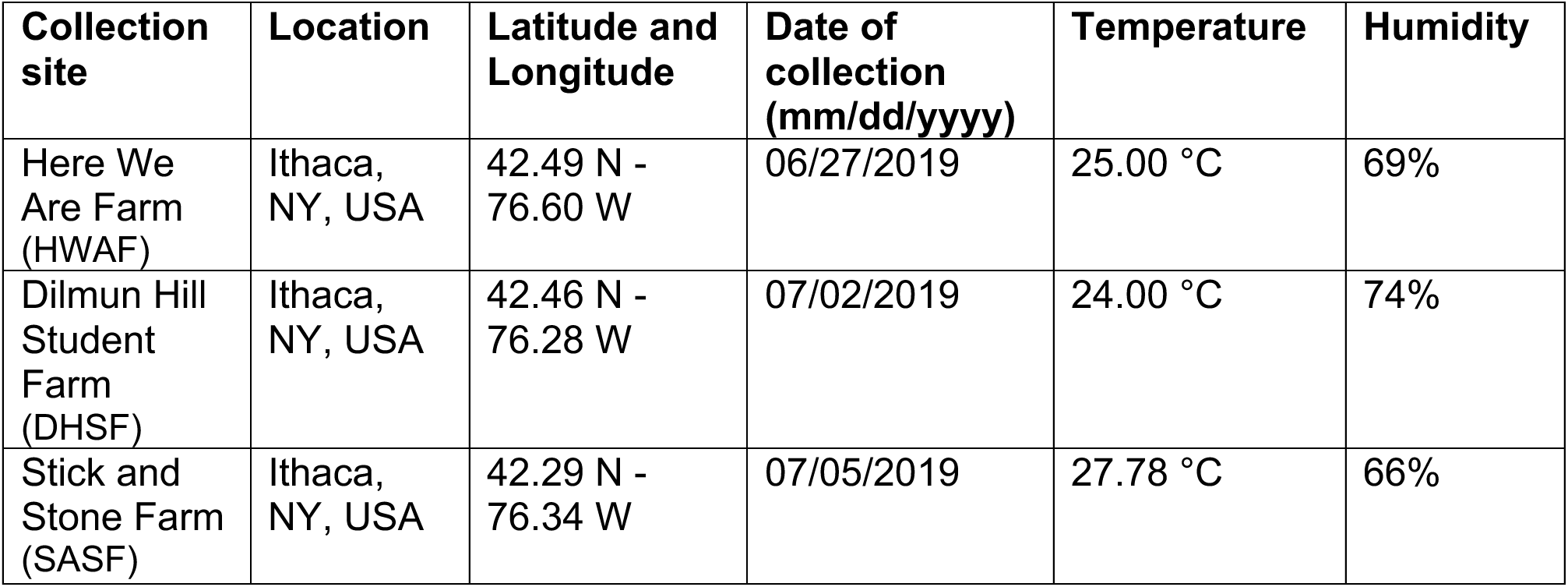
Locations and environmental conditions of phyllosphere sample collection.

To limit bias caused by potential aphid preferences, we sampled plants that had feeding pea aphids and those that did not. At each location we chose nine plants, three with aphids and six without, spaced approximately 1-5 meters apart. From plants with aphids, we collected three leaf pairs with aphids as well as three leaf pairs without aphids, distributed as evenly as possible across the plant. From plants without aphids, three leaf pairs were collected. This resulted in 36 samples per location: nine leaves with aphids and 27 without, and 108 samples in total. Leaves were collected in individual sterile 50mL conical tubes using scissors sterilized with 70% EtOH. Tubes were kept in a cooler with ice packs during collection and transport.

Immediately after transport to the lab aphids were removed from leaves using a sterilized paint brush. Leaf samples were cut into smaller pieces using sterilized scissors and 10 mL of sterile 10 mM MgCl_2_ buffer was added. Samples were sonicated for 10 minutes to dislodge epiphytic bacteria from the leaf surface and then briefly vortexed. To remove plant cells prior to DNA extraction, the leaf wash was passed through a 5 µm filter. Fractions of the supernatant were used for DNA extraction and culture isolation.

### Sequencing procedures

To investigate the diversity and functional potential of fluorescent isolates from pea plants, we sequenced genomes of isolates recovered from each of our field sites. To do so, 50 µL of each leaf wash was spread onto King’s B (KB) agar plates supplemented with the antifungal nystatin (35 ug/ml). Plates were incubated at 27°C for 24 hours, and then fluorescent colonies were identified using a Wood’s UV flashlight. Isolates selected for genome sequencing represented the morphological diversity of fluorescent colonies on each plate, with 0 - 3 isolates per plated sample selected. Out of 108 samples, 36 yielded at least one fluorescent isolate, resulting in 48 isolates in total. Fluorescent isolates were grown overnight in liquid KB media and cells were harvested for DNA extraction using a Qiagen DNeasy blood and tissue kit following the recommended protocol for cultured cells. Sequencing was performed at the Cornell Biotechnology Resource Center with Nextera WGS library preparation and Illumina NextSeq500 paired-end 2 x 150 bp reads.

To assess total bacterial communities on pea plants and the prevalence of pseudomonads at our field sites, we used the remaining supernatant for metagenomic sequencing. Samples were centrifuged (3650 RCF for 10 minutes) to pellet bacterial cells and DNA was extracted using a Qiagen DNAeasy Blood and Tissue kit with one modification. We included a bead beating step following the addition of lysis buffer to ensure efficient mechanical lysis of both Gram-positive and Gram-negative bacterial cells. Silicon microbeads (0.4 mm) were added to each sample prior to beating in an MP-bio Fastprep 24 beadbeater at 4.0 M/S for 15 seconds. Metagenomic sequencing was performed at the Cornell Biotechnology Resource Center with Nextera WGS library preparation and Illumina NextSeq500 paired-end 2 x 150 bp reads.

### Sequence analysis

Of 108 samples collected, 84 yielded sufficient sequencing reads for downstream metagenomic community analysis. Raw metagenomic reads contained on average 2,716,250 read pairs per sample. After quality control using FastQC (Wingett and Andrews 2018) and adaptor trimming with Trimmomatic (Bolger et al. 2014), reads were filtered using BWA-MEM (Li and Durbin 2010; Li 2013 Mar 16) with reference genomes for pea aphid (GenBank accession NC_042493) and snap pea (PUCA010000002). The resulting filtered sequences averaged 385,439 reads per sample (Supplementary Table S1). Using default settings throughout, reads were assembled with metaSPAdes (SPAdes v. 3.15.2) (Bankevich et al. 2012; Nurk et al. 2017), binned with Metabat2 (Kang et al. 2019) to isolate distinct metagenomic assembled genomes (MAGs), and the resulting bins were checked for contamination and completeness using CheckM (Parks et al. 2015) (Supplementary Table S1). MAGs with >80% completeness and <10% contamination were identified with species-level genome bins (SGBs) from metagenomes (MetaPhlAn v. 4.0) (Blanco-Miguez et al. 2022) (Supplementary Figure S2).

We used the phyloFlash pipeline (Gruber-Vodicka et al. 2020) to determine the taxonomic composition of phyllosphere communities. In this pipeline, metagenomic reads were mapped to the SILVA 138 database (SILVA 138.1 SSRef NR99) (Quast et al. 2013) and reads with >70% identity were used to assemble full-length small-subunit (SSU) rRNA gene sequences with SPAdes (Bankevich et al. 2012). Taxonomic affiliation was determined according to the reference sequence to which the SSU rRNA reads mapped, with reference sequences clustered by 97% identity. This output was filtered to include only 16S, not 18S rRNA gene sequences. Since these units are not strictly the same as the commonly used Operational Taxonomic Units (OTUs) or Amplicon Sequence Variants (ASVs), we refer here to taxonomic units, or simply taxa.

Of the 84 metagenomes, 82 yielded at least one full-length SSU rRNA sequence assigned via the phyloFlash pipeline using default settings. Samples were filtered to include those with more than one SSU rRNA assignment, and communities were filtered to exclude eukaryotic and unassigned taxa. Out of 84 total samples, 76 passed these quality control metrics and were used for downstream analysis.

To confirm the prevalence of pseudomonads and pyoverdine biosynthesis potential in phyllosphere metagenomic communities, we aligned reads from our metagenomes to the *Pseudomonas syringae* pv. *syringae* strain B728a (GenBank assembly NC_007005.1) housekeeping marker gene *gyrB* (PSYR_RS00020, NC_007005.1:3809-6226) (Yamamoto et al. 2000; Gomila et al. 2015) and the pyoverdine biosynthesis gene *pvdL* (PSYR_RS10105, NC_007005.1: 2217771-2230781) (Bultreys et al. 2001; Feil et al. 2005). Alignment was performed using Bowtie2 (Langmead and Salzberg 2012) with default settings. To directly compare alignment between these references with different lengths, the number of aligned reads for *pvdL* were normalized to the total sample reads multiplied by the length ratio between *gyrB* and *pvdL* (5.38).

From sequencing of cultured isolates, raw genomic reads contained an average of 1,188,340 read pairs, with an average length of 176,297,883 bp. Genomes were assembled with SPAdes v 3.15.2 utilizing the default isolate settings. Assemblies were checked for contamination and completeness with CheckM and QUAST (Gurevich et al. 2013). Phylogenetic analysis was performed using the PhyloPhlAn 3.0.51 pipeline for amino acid sequences (Asnicar et al. 2020) including default settings for DIAMOND, MAFFT, trimAI, and RAxML for the configuration, and the medium diversity and accurate options selected for the run settings (Stamatakis 2006; Capella-Gutiérrez et al. 2009; Katoh and Standley 2013; Buchfink et al. 2015). The best-fitting substitution model according to ModelFinder was VT+F+R3 (Kalyaanamoorthy et al. 2017). Identity of the assembled genomes (isolates and SGB quality MAGs) was determined using JSpeciesWS (Richter et al. 2016). Using JSpeciesWS, closely related reference genomes were selected from top hits of a tetra-nucleotide Correlation Search (TCS) of assembled genomes against the curated Ensembl reference database Genomes DB. Assembled genomes and references with higher than 95% calculated Average Nucleotide Identities (ANI) were considered identical at the species level.

The diversity of siderophore biosynthesis and receptor genes, as well as biosynthesis genes for the fluorescent compound phenazine, in phyllosphere isolates was determined through genome annotation with RAST (Aziz et al. 2008). Annotations for genes of interest were confirmed using NCBI BLAST (Altschul et al. 1990; Sayers et al. 2021) with known ortholog sequences as queries (Supplementary Table S2).

### Fluorescence quantification

To quantify the fluorescence spectra of isolates in standard culture conditions, bacterial lawns of each strain were imaged under UV light (60 W fluorescent blacklight bulb, Adkins Professional lighting). Lawns of each isolate were prepared by inoculating petri dishes of KB agar media with liquid cultures grown overnight. To ensure uniform lawn growth, 3mL of overnight culture was poured onto the solid media and left to soak for 30 minutes. Excess liquid was then poured off and the plate was left to dry for 10 minutes. Pictures of the plates were taken following a 24-hour incubation period at 28°C. A spectrally flat polytetrafluoroethylene surface was placed under each plate to uniformly reflect light (Hendr et al. 2018). Color and 18% grey standards were used as controls for each photo. Pictures were taken as RAW images (a file format that provides unprocessed and objective image data) using a Canon am IS00 camera set to ISO of 500. To quantify both fluorescence color and UV reflectance of strains, images were taken with and without a UV lens filter.

For analysis, RAW images were imported into ImageJ and converted into multispectral image files using Sensory Ecology’s micaToolbox plugin (Troscianko and Stevens 2015). These multispec images were made by combining the visible light photos with UV photos both standardized to a 18% Grey standard. Multispec images were then converted to .tiff files and a circular region of interest of the same size (50 mm diameter) was selected for each photo. Tiff files were then run through a custom macro (Miller et al. 2019) to randomly sample 1000 subregions and record values of each subregion within three binary color channels (vRed, vBlue, vGreen) as well as two UV sensitive channels (UVR and UVB). Color values for each channel are then recorded for each of the 1000 randomly selected regions of interest and exported into Excel. This data was then combined and imported into R and analyzed using plotly and ggplot2 packages (Wickham 2016; Sievert 2020).

### Aphid behavior experiments

To determine if there is a relationship between fluorescent emission of phyllosphere isolates and aphid behavior, four pairs of isolates were chosen based on their fluorescent color and intensity and tested for aphid behavioral responses. Based on previous results, we hypothesized that strains with more blue fluorescence would be avoided. We therefore chose pairs of strains with more blue or more green biased fluorescence from our measurements. More intensely fluorescent strains may elicit a stronger response. They would also absorb more UV light and thus decrease UV reflectance. Isolates were therefore selected such that each pair containing a blue and a green biased isolate had similar UV reflectance values, with pairs selected from across the range of fluorescent intensities. We also included a positive control strain previously shown to deter aphids in these experiments (Hendry et al. 2018).

Our experiment sought to measure the preference of winged dispersing (alate) aphids settling on plants colonized epiphytically with fluorescent strains versus control treated plants. Fava bean plants (*Vicia faba*) were sprayed with either a bacterial suspension in 10 mM MgCl2 (buffer) or a control of buffer alone. Cultures were grown in KB media at 28°C overnight and were centrifuged at 3650 RCF for 10 minutes to pellet cells. Pellets were washed twice via resuspension with 10mL buffer before adjusting the bacterial suspension to an OD_600_ of 1.0 (+/- 0.02). Bacterial suspensions were sprayed onto two-week-old fava bean plants using a sterilized spray bottle (Smee and Hendry 2022). Plants were sprayed until visibly wet but not dripping (about 3 sprays per spray area) and left to dry for one hour in a biosafety cabinet. Once dry, plants were placed in a 180 x 71 x 71 cm greenhouse tent kept at 21°C and ∼80% relative humidity. Full spectrum T4 cool blue grow lights were suspended ∼40 cm above each plant. Supplemental UV light was provided in the form of two 60 W fluorescent blacklight bulbs (Adkins Professional lighting) per tent. We previously found this set up to be more similar to natural sunlight than full spectrum bulbs alone (Hendry et al. 2018). Lights were kept on a 16:8 hour day night cycle.

Pea aphid clone CWR09/18 was used for experiments. This line was collected in 2009 by Angela Douglas in Freeville, NY and is free of all endosymbionts except the obligate *Buchnera aphidicola* (Bouvaine et al. 2012; Macdonald et al. 2012). Lab colonies were reared at 20°C under 16:8 hr light cycle and fed on fava bean. Alate aphids were generated by allowing cages to become overpopulated until winged individuals were attempting to disperse. Winged dispersing aphids have been shown to have different color preferences than other life stages (Döring and Chittka 2007), and represent an ecologically realistic stage to be choosing among host plants. Alates were introduced to each greenhouse tent via petri dishes at two points equally spaced apart, hung 20 cm above plants. As such, the aphids were forced to fly to plants and could choose to land on plants treated with either the bacterial suspension or the buffer. Each tent contained 12 plants (6 of each treatment) arranged in a matrix with ∼30 cm between each plant. Plants in each replicate were randomly assigned positions in this matrix. About 30-50 alates were placed in each petri dish. Assays were replicated five times for each isolate except for the positive control strain B728a, which was replicated three times. After 24 hours, the number of aphids which had settled on each plant were counted.

Pea aphids typically feed on the undersides of leaves or apical growing tips of plants. They can be difficult to accurately count on apical tips where they lodge between leaf buds and fall off plants easily when disturbed. Additionally, actively growing leaf buds are likely to have received a lower bacterial inoculum than fully grown leaves. We therefore limited plant disturbances by counting aphids which settled onto fully grown leaves or were visible on stems. We confirmed bacterial growth on leaves by measuring colony forming units (CFUs) from 10 leaf disks sampled from across each plant. Leaf disks were placed in 10mL of 10 mM MgCl2 buffer and sonicated for 10 minutes, then briefly vortexed to release epiphytic bacteria. Serial dilutions of this suspension were plated onto KB media supplemented with the antifungal nystatin. Plates were incubated at room temperature for 48 hours before fluorescent colonies (identified with a Wood’s lamp) were counted. On average CFUs from 10 leaf disks ranged from 3.28 x 10^2^ to 3.93 x 10^5^ CFUs per leaf disk area (2.58 cm^2^), which are ecologically relevant densities (Smee and Hendry 2022).

### Statistical analysis

Microbial community analysis was performed in R version 4.3.2 (R Core Team 2023), using the phyloseq and vegan packages (McMurdie and Holmes 2013; Oksanen et al. 2022). The species diversity (alpha diversity) of phyllosphere metagenomic communities across distinct variables was evaluated using Shannon diversity and Observed richness metrics and compared using pairwise Wilcoxon Rank Sum tests. We found diversity did not vary significantly between locations, host plant, sample type, or sample leaf.

Community compositions (beta diversity) were evaluated using Non-metric Multidimensional Scaling (NMDS) of Bray Curtis dissimilarities. We did not find significant clustering for any of the variables tested: location, host plant, or sample type (Supplemental Figure S1).

To determine community level differences between phyllosphere metagenomes, we employed Permutational Multivariate Analysis of Variances (PERMANOVA). PERMANOVAs were performed in R using the vegan (v.2.6-4) and ade4 (v.1.7-22) packages (Dray and Dufour 2007; Oksanen et al. 2022). The number of permutations specified was always 999. To examine community level variations across sample locations, we used a PERMANOVA with Bray Curtis metrics, with individual host plant included as a nested variable (formula: ‘dist ∼ Location/PlantID’). To further determine the impact of each location on community variation, we performed pairwise-PERMANOVAs on community compositions between locations (Supplemental Table S5).

Correlations that did not involve microbial communities were evaluated using Spearman Rank Correlation Tests in R.

Aphid preference was recorded as a percentage, calculated from the number of aphids settled on control plants over the total settled aphids counted per replicate tent (sum of aphids on control plants divided by sum of total aphids observed). For this analysis, our null expectation is that 50% of aphids should choose to settle on control treated plants. We define altered aphid behavior to mean that significantly more or less than 50% of aphids chose control plants over the treated plants. All statistical analyses were conducted in R (v4.3.0). We used binomial generalized linear mixed-effects models (GLMMs) with the R package lme4 (Bates et al. 2015). To test for deviations from the null expectation we included -1 as a variable in GLMMs, effectively removing the model intercept. The resulting p value indicates if the probability of aphids choosing control plants significantly varied from 50%. Date and experimental replicate were included as random effects. Strain identity was added to the null model as an explanatory variable, and likelihood ratio tests were used to determine whether strain identity significantly affected aphid choice compared to the null expectation. The package ‘emmeans’ was used to obtain probabilities with confidence intervals from the models (Lenth 2024).

## Results

### P. sativum epiphytic bacterial community composition

To our knowledge, the bacterial composition of the pea plant phyllosphere has not been previously assessed. To determine the prevalence and relative abundance of fluorescent pseudomonads within the context of *P. sativum* phyllosphere microbiomes we therefore assessed total epiphytic bacterial communities from pea leaves at three sites. Using metagenomic sequencing we determined the relative abundance of full length16S rRNA gene sequences obtained from 76 leaf washes. We found a consistent cohort of bacteria; by descending prevalence across samples common bacterial families were *Flavobacteriaceae*, *Erwiniaceae*, *Pseudomonadaceae*, *Enterobacteriaceae*, *Comamonadaceae*, *Staphylococcaceae*, and *Crocinitomicaceae* (Figure 1). These taxa are prevalent in samples across all locations sampled and are consistent with common phyllosphere community members found in other systems (Dong et al. 2019; Bashir et al. 2022). Our results did not find a high relative abundance of other taxa that are often found in phyllosphere communities, notably taxa in the orders Bacillales, Hyphomicrobiales, Sphingomonadales, and Micrococcales (Morella et al. 2019; Bashir et al. 2022; Mechan-Llontop et al. 2023; Mehlferber et al. 2023). Our sequencing depth was targeted to abundant taxa to best assess common pseudomonads and may have been insufficient to recover all taxa present in these communities.

**Figure 1:**
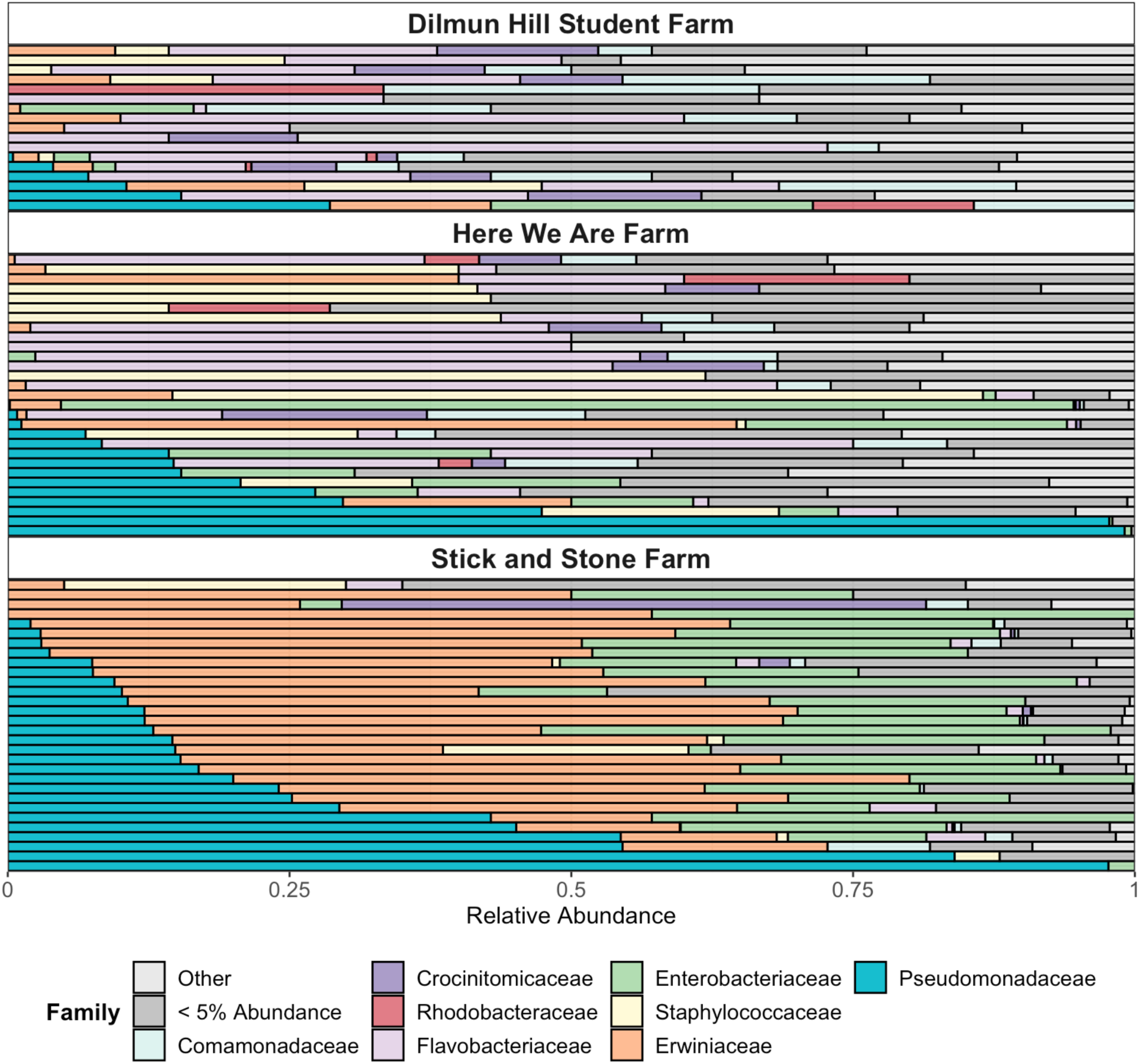
Relative abundance of bacteria in epiphytic communities from *P. sativum* leaves. Each row represents a single leaf sample with sample location given. Taxonomic assignments, shown here are at the family level, were assigned to full length assembled SSU rRNA sequences mapped to reference sequences (clustered with 97% identity) from the SILVA 138.1 database. Taxa with a relative abundance of <0.1% were not included.

Location was significantly correlated with bacterial community composition (Supplementary Table S5: pairwise PERMANOVA, p<0.05 for all location comparisons), but not alpha diversity (Observed richness, p >0.05 and Shannon Index, p >0.05). Within a site, identity of sampled plant was significantly correlated with phyllosphere microbial community composition at HWAF (p= 0.019) and SASF (p=0.018), but not DHSF (p=0.43). Overall, we find that location accounted for 20% of variation in community composition (PERMANOVA, R2 = 0.203, p = 0.005), and individual host plant accounts for 30% (R2 = 0.304, p = 0.005). We did not find any significant correlation of aphid presence with community composition (PERMANOVA, R2 = 0.83 p = 0.397).

Samples from SASF were distinct from other sites due to having a higher prevalence of gammaproteobacteria (Erwiniaceae, Enterobacteriaceae, and Pseudomonadaceae, 93%, 90%, and 86% of samples, respectively) compared to other sites, where the highest prevalence of these was Pseudomonadaceae at HWAF (48% of samples). In comparison, Flavobacteriaceae were prevalent in samples from DHSF and HWAF (88% and 76% of samples), but less common in samples from SASF (45% of samples) (Figure 1). We note that sampling at SASF occurred under warmer conditions (27.78°C), which could explain the abundance of fast growing gammaproteobacterial taxa (Aydogan et al. 2018).

### Prevalence of fluorescent pseudomonads in the P. sativum phyllosphere

Pseudomonads were common across all samples and sites. In total, 60.5% (46/76) of *P. sativum* epiphytic samples contained taxa from the Pseudomonadaceae (Figure 1). When present, the average relative abundance of Pseudomonadaceae within samples was 23.5%, second only to Erwiniaceae (28.95%). In nine samples, Pseudomonadaceae was the most abundant family, with relative abundance as high as 99.08%. Despite this, we did not find a correlation between Pseudomonadaceae abundance and sampling location (PERMANOVA, R2 = 0.041, p = 0.841), host plant individual (PERMANOVA, R2 = 0.435, p = 0.787), or the presence of aphids at the time of sampling (PERMANOVA, R2 = 0.021, p = 0.444). Pseudomonad diversity in these samples was also high; 16S rRNA gene sequences from a total of 104 *Pseudomonas* taxa (species and pathovars) were identified across all *P. sativum* phyllosphere communities. Furthermore, communities that contained co-occurring pseudomonad taxa were common. Of samples containing *Pseudomonas* identified at the species level, most (70%, 26 out of 37) were found to harbor multiple *Pseudomonas* species, with a majority (65%, 24 out of 37) housing at least three co-occurring species.

Genomic assembly from across 76 samples yielded sixteen MAGs with more than 60% completeness, all of which belonged to the genera *Pantoea* and *Pseudomonas*. This is consistent with the abundance of Erwiniaceae and Pseudomonadaceae 16S rRNA gene sequences. Four species were identified among the MAGs: *Pantoea agglomerans*, *Pantoea eucrina*, *Pseudomonas putida*, and *Pseudomonas syringae*. Fifteen phyllosphere samples yielded a single MAG each, with one sample yielding two (Supplemental Figure S2). *Pantoea* and *Pseudomonas* are both known to be common epiphytes (Vorholt 2012; Morris et al. 2013; Baltrus et al. 2014; Flury et al. 2016; Bashir et al. 2022) and our ability to recover MAGs from them here, as well as the 16S rRNA gene analysis above, supports the conclusion that these taxa were abundant in the epiphytic microbial communities of *P. sativum*.

To determine the prevalence of potential pseudomonad fluorescence within *P. sativum* epiphytic bacterial communities, we searched our metagenomic sequence data for a gene underlying biosynthesis of the fluorescent siderophore pyoverdine, *pvdL* (Mossialos et al. 2002; Schalk et al. 2020). We found that the potential to synthesize pyoverdine was relatively common in epiphytic communities (Figure 2). Reads from 59% of samples (45/76) aligned to *pvdL* at least once. The percentage of reads per sample that mapped to *pvdL* was similar to percentages of mapping to a *Pseudomonas gyrB* sequence, a housekeeping gene that we used as a proxy for pseudomonad abundance, and also to pseudomonad abundance from 16S rRNA gene sequences (Figure 2). For instance, 81% of samples with alignments to *gyrB* also align to *pvdL*, suggesting an infrequent occurrence of pseudomonads unable to synthesize pyoverdine. These data show that most of the pseudomonads in our samples have the potential to be fluorescent and impact aphid behavior due to pyoverdine production.

**Figure 2:**
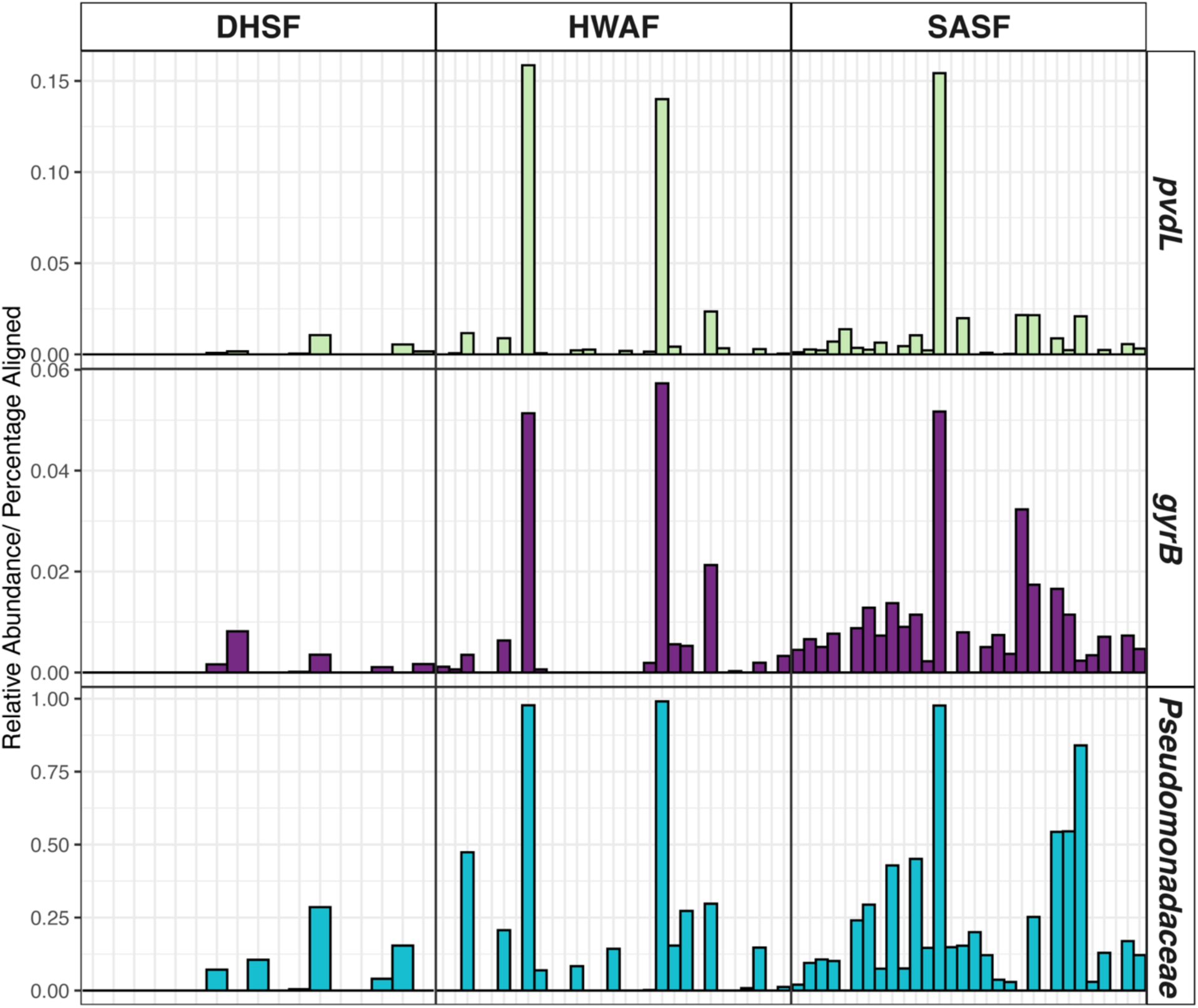
Normalized percentage of reads in epiphytic metagenomic samples (bottom axis) aligned to the pyoverdine biosynthesis gene *pvdL* (top row, green bars) and housekeeping gene *gyrB* (middle row, purple) from *P. syringae* pv. *syringae* B728a. To account for differences in gene length, alignment values for *pvdL* were normalized to the total sample reads * the length ratio between *gyrB* and *pvdL* genes (5.38). Relative abundance of reconstructed *Pseudomonadaceae* 16S rRNA gene sequences in metagenomes are included for comparison (bottom row, cyan bars).

### Genetic and fluorescence diversity of pseudomonad isolates

Sequenced genomes from fluorescent pseudomonad isolates belonged to diverse lineages (Figure 3). Isolates were spread across the species complexes of *Pseudomonas putida*, *Pseudomonas syringae*, and *Pseudomonas fluorescens* (Berge et al. 2014; Garrido-Sanz et al. 2016; Lalucat et al. 2020; Babalola et al. 2021; Girard et al. 2021). We performed average nucleotide identity (ANI) analysis on a subset of isolates that were used in behavior experiments. Based on highest ANI similarities and the species threshold of 95%, we determined that these eight isolates belong to seven species from these species complexes (Supplementary Table S3).

**Figure 3:**
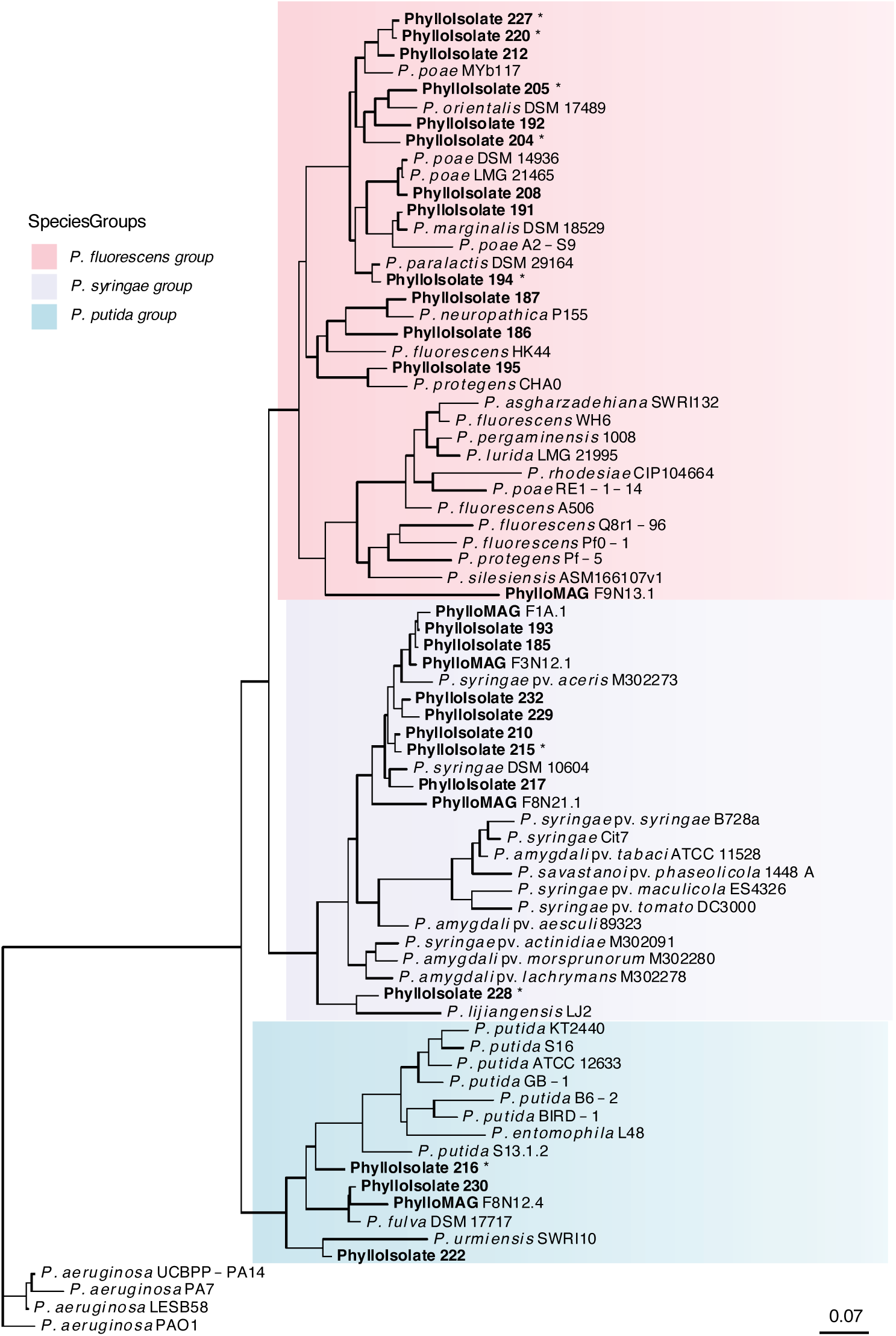
Whole genome maximum likelihood tree of fluorescent epiphyte isolates and related *Pseudomonas*. Isolates and metagenomic assembled genomes from this study are bolded. Other representative genomes for diverse *Pseudomonas* species were obtained from GenBank and chosen based on average nucleotide identity to isolates. Background colors indicate *Pseudomonas* species complex. Bootstrap values less than 0.9 are shown. Asterisks indicate isolates used for choice assays.

We investigated the functional fluorescence diversity of the pseudomonad isolates collected here plus some used in previous work (Hendry et al. 2018), 56 isolates in total. We focused on the color of fluorescence from these isolates, as well as the intensity of their fluorescence, as we had previously found that the intensity of blue fluorescence correlated with aphid avoidance (Hendry et al. 2018). We found high variation across isolates grown in standard culture condition for both the intensity of fluorescence color, as well as how biased towards blue versus green the fluorescence color was (Figure 4A). Blue and green color values span a full range from very low for media alone and a pyoverdine deficient mutant (ΔpvdL), to faintly fluorescent isolates such as the previously studied *P. syringae* pv. *aptata* (Hendry et al. 2018), through to brightly blue and/or green fluorescent isolates (Figure 4B). The intensity of these color values correlated negatively with measured values of UV reflectance (Supplementary Figure S3), demonstrating that higher amounts of absorbed UV light, and therefore lower amounts of UV reflectance, can be used as a proxy for fluorescence intensity. Strains that we previously found to be avoided by aphids (*P. syringae* pv. *syringae* strain B728a, *P. syringae* strain Cit7, and *P. syringae* pv. *japonica*) had moderately intense fluorescence that was average to moderately blue biased (Figure 4) (Hendry et al. 2018). The fluorescence measures of most newly collected strains were more intense and often more blue or green biased. Epiphytic pseudomonads therefore have the potential to produce fluorescence that covers a broad range of intensities and colors.

**Figure 4:**
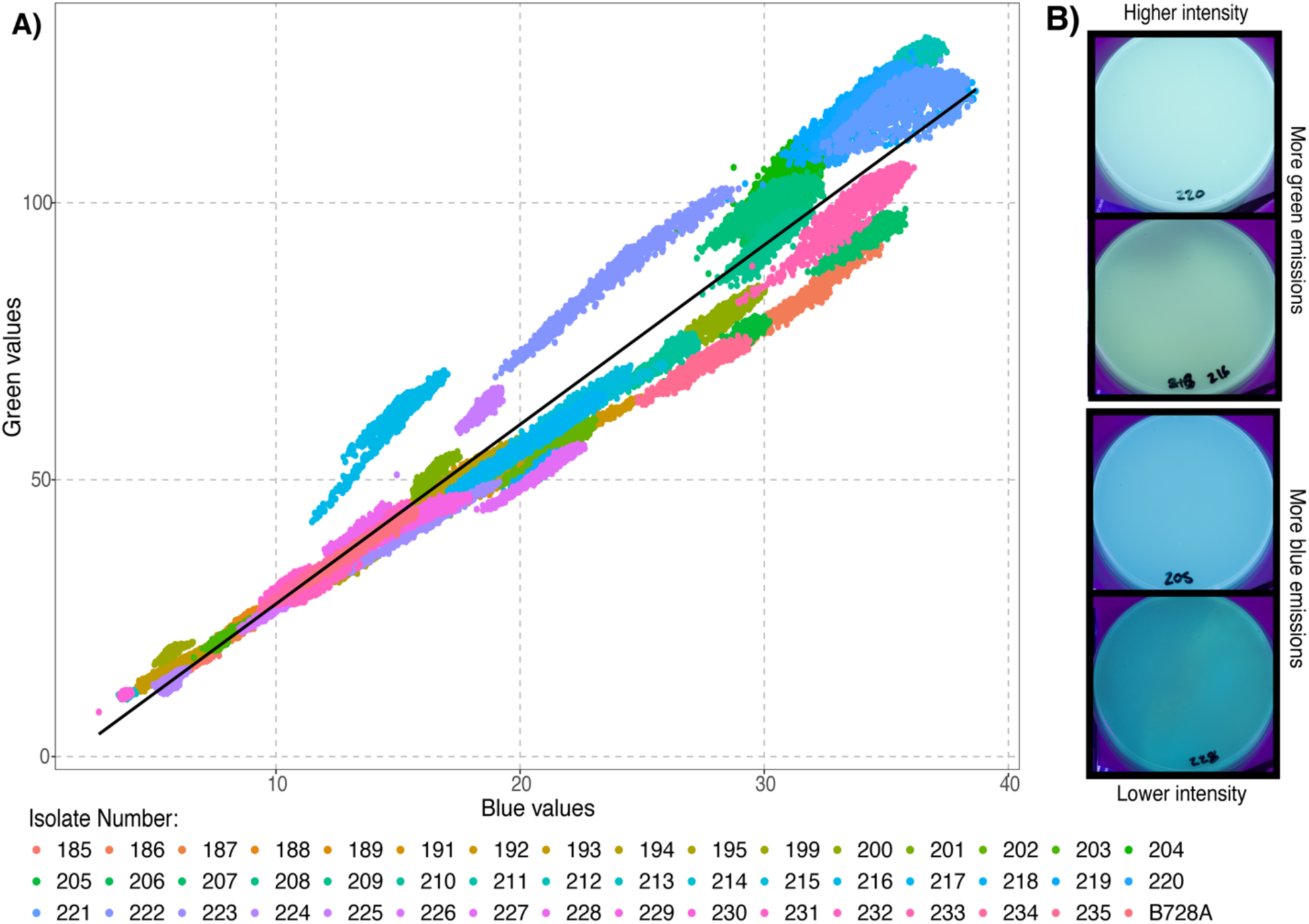
**A)** Blue and green color intensity from fluorescent epiphytic *Pseudomonas* isolates grown in standardized culture conditions on agar plates and photographed under ultraviolet light. A linear regression model best fit line is shown. Each point represents the blue and green values from a single pixel in the plate image. Points are colored by isolate identity. **B)** Example isolate plates photographed under UV light. Representative strains with fluorescence that was higher and lower intensity, as well as more green versus blue biased, were chosen. Raw, uncorrected and unprocessed images are shown.

This diversity of fluorescence could be due to variation in pyoverdine structure across isolates, or it could be influenced by the production of additional siderophores or fluorescent compounds. Sequenced genomes from our isolates show variation in repertoires of siderophore synthesis genes (Figure 5). We recovered pyoverdine synthesis and receptor genes in most isolate genomes. Genes to produce and bind the siderophore achromobactin (Berti and Thomas 2009; Owen and Ackerley 2011) were also fairly common. One isolate separately had biosynthesis and receptor genes for yersiniabactin (#228) (Bultreys et al. 2006; Jones et al. 2007) while several isolates had receptor genes for aerobactin (#185, #195, #217, #222, and #230) (de Sousa et al. 2022; Schalk and Perraud 2023). Achromobactin, yersiniabactin, and aerobactin are not reported to be fluorescent. However, we did recover biosynthesis genes in two strains for pyochelin, a *Pseudomonas* secondary siderophore which is fluorescent (Figure 5) (Namiranian et al. 1997; Youard et al. 2007; Andrić et al. 2023). We additionally found that several strains have a biosynthesis gene for phenazine (Figure 5), a fluorescent pigment that is common in pseudomonads (Mavrodi et al. 2006; Wang et al. 2011; Mavrodi et al. 2013; Vilaplana and Marco 2020). Production of these molecules could contribute to fluorescence color.

**Figure 5:**
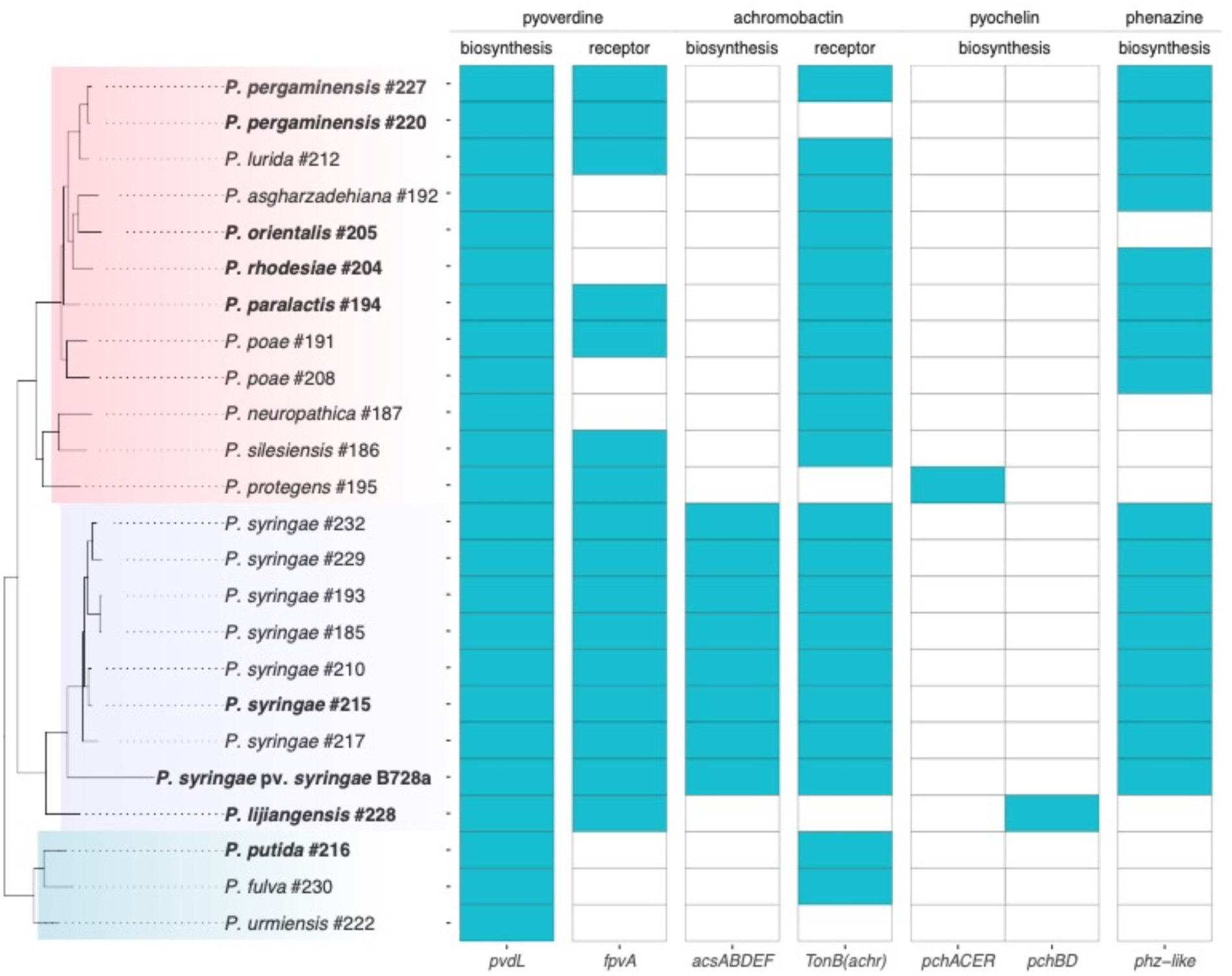
Biosynthesis and receptor genes for common siderophores and fluorescent molecules identified in the genomes of fluorescent *Pseudomonas* isolates used for choice assays (bold) from this study. Boxes indicate if gene is found in the respective genome (cyan) or not (white). Gene names are listed in the bottom axis, with gene function and siderophore on the top axis. Phylogeny of sequenced isolates is adapted from Figure 3.

### Aphid behavioral responses to diverse fluorescent epiphytes

We performed behavioral experiments on a subset of nine strains with variable fluorescence color and intensity to determine how fluorescent colors of pseudomonads influence aphid feeding choice. We found that these epiphytic strains caused a diversity of behavioral responses in dispersing alate aphids. Consistent with previous work, we found that *P. syringae* pv. *syringae* strain B728a was avoided by aphids (Figure 6A, Supplementary Table S4). We found an additional two isolates (#215 and #204) which were also avoided, slightly less on average than *P. syringae* pv. *syringae* strain B728a. An additional four isolates elicited no significant behavioral response. Although two of these showed an average tendency to cause avoidance (#205 and $194), this was not significantly different than chance.

**Figure 6:**
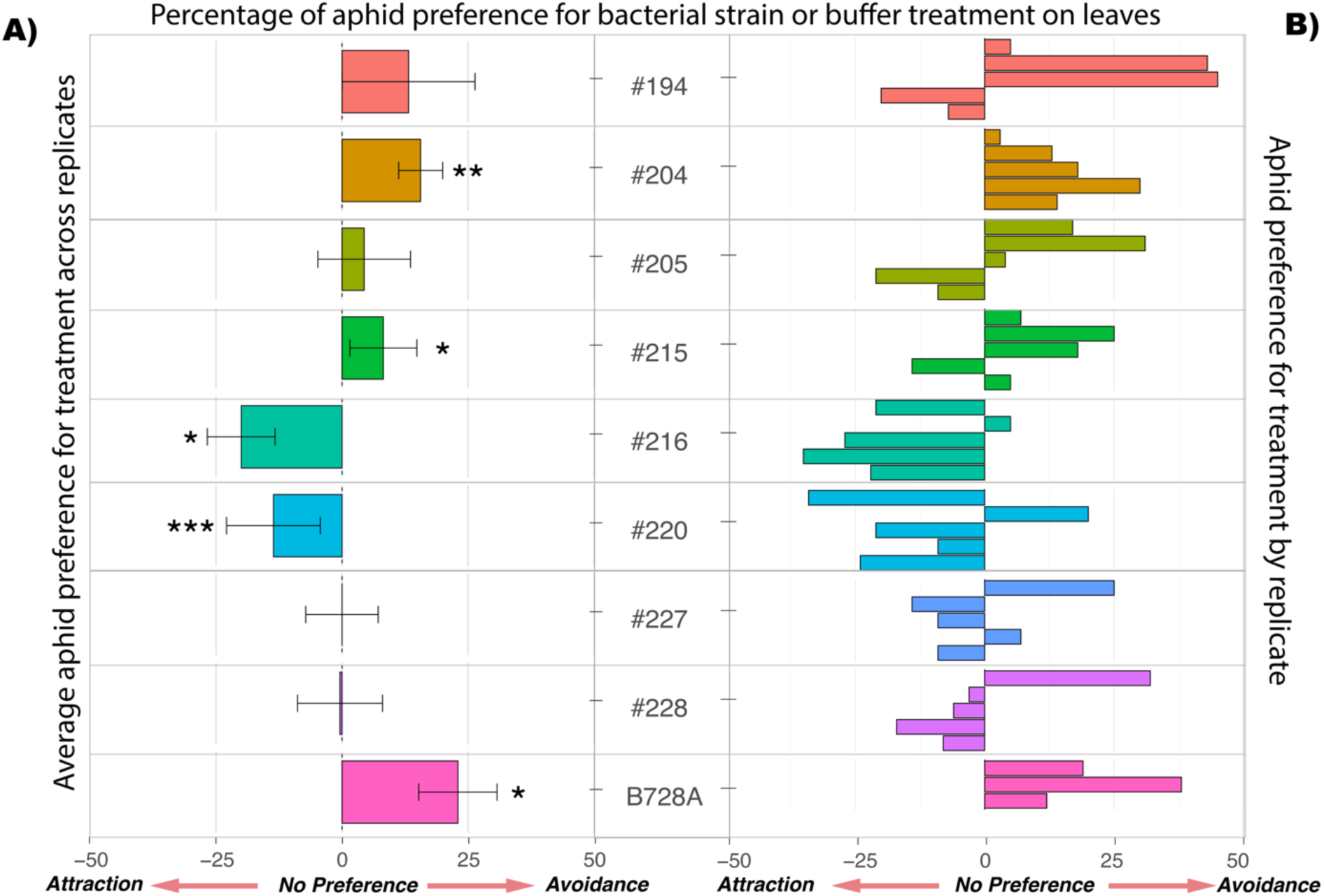
Mean aphid choice for plants treated with selected isolates versus control treated plants (A) and histogram of aphid choice across replicate blocks for each isolate (B). Aphid choice is shown as the percentage of aphids settled on control treated plants out of all settled aphids, relative to 50% of aphids on control treated plants. Positive values therefore indicate avoidance of bacterially treated plants, while negative values indicate attraction to bacterially treated plants.

Surprisingly, plants with epiphytic populations of two isolates (#216 and #220) were significantly preferred by aphids over control treated plants (Figure 6A, Supplementary Table S4). When these isolates were present on plants an average of 64% (#220) and 70% (#216) of aphids settled to feed on bacterially treated plants rather than control plants. This suggests that epiphytic bacteria may have previously unappreciated effects on aphid populations: in addition to deterring aphids they could also encourage aphid settling and feeding.

Overall, we find a poor correlation between our measurements of fluorescence color in culture and the effects of a given isolate on aphid behavior (Figure 7). Furthermore, we did not find that measures of UV reflectance in culture correlated with behavior (Spearman Rank Correlation, rho = 0.31, p = 0.42) suggesting that aphid detection of UV light was not influencing preferences directly. We also found no pattern between the siderophore repertoire of an isolate’s genome (Figure 5) and the behavioral response that aphids show to a isolate on plants (Figure 6:Spearman Rank Correlation, rho = 0.11, p = 0.42). Siderophores are typically not produced under all environmental conditions, and therefore the potential to produce a siderophore does not necessarily equate to actual production on plants. We found high variation in aphid responses across replicate experiments for some isolates (Figure 6B). For instance, isolate #205 was avoided in some replicates, with up to 81.5% of aphids on control plants, and attractive in other replicates with as few as 29.4% of aphids choosing control plants. This suggests that the fluorescence produced by an isolate may vary across replicate experiments. Variation in siderophore production can be driven by environmental conditions such as temperature and nutrient availability (Garibaldi 1971; Duffy and Défago 1999; Katiyar and Goel 2004; Vindeirinho et al. 2021). Consistent with this, we found that the green versus blue bias of fluorescence from isolate #194 in culture changed depending on the carbon source available (Figure S4). Environmental variability on plants could influence variation in aphid behavior across replicates.

**Figure 7.**
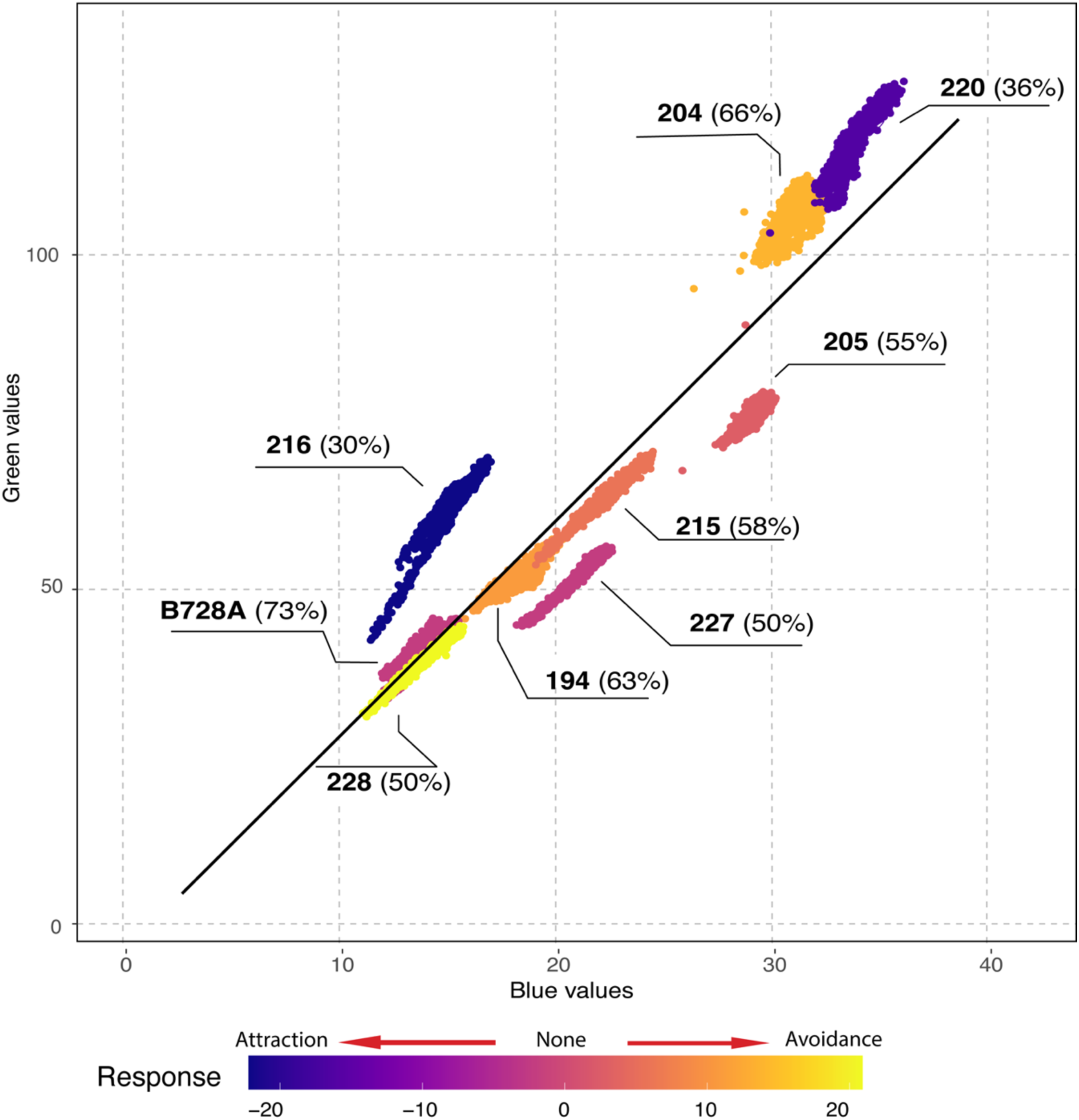
Fluorescence color and intensity values (data from figure 4), as well as best fit line, for isolates used in behavioral assays. Points are colored along a scale to indicate aphid behavioral response using the mean aphid choice values across replicates for each strain. Yellow colors indicate strains that are more avoided by aphids, purple colors indicate strains that are more attractive to aphids, and pink colors indicate no preference. Color gradient is calculated using the proportion of aphids on control plants, over or under 50%. The absolute proportion of aphids on control plants is given in parentheses by each strain name.

## Discussion

We identified a cohort of abundant bacteria in the *Pisum* phyllosphere community, which were prevalent in the metagenomes of sampled leaf washes. Relatively consistent, plant specific phyllosphere communities have also been found associated with other plant hosts (Aydogan et al. 2018; Dong et al. 2019; Dong et al. 2019; Bashir et al. 2022; Meyer et al. 2022; Debray et al. 2023; Mechan-Llontop et al. 2023). Individual host plant identity and location were the factors with the most influence on variation in community composition and relative abundance of bacteria on *Pisum* leaves. For example, SASF had higher relative abundances of *Pseudomonadeae* and other gammaproteobacteria, compared to Flavobacteriaceae, which were common at other sites. This location was sampled on a warmer day, similar to the optimal growth temperature for certain *Pseudomonas* taxa (Tribelli and López 2022). This variation between locations could be due to differences in environmental conditions such as temperature, humidity, and UV exposure from sunlight (Vorholt 2012; Aydogan et al. 2018; Legein et al. 2020; Chai et al. 2023; Özkaya et al. 2023). Other factors, such as differences in soil composition, individual host plant physiology, and exposure to other organisms (insects and humans) could explain variation across individual plants (Humphrey et al. 2014; Granzow et al. 2017; Chartrel et al. 2021; Meyer et al. 2022). Overall, we find the phyllosphere community of *Pisum* leaves to be relatively consistent for abundant taxa.

We found that fluorescent pseudomonads with the potential to impact aphid behavior were relatively common epiphytes in *P. sativum* phyllosphere communities. Pseudomonads were found in the majority of phyllosphere samples, and were also among the most prevalent epiphytes in phyllosphere communities within each location. Pseudomonads are known to disperse via the water cycle (Morris et al. 2013), which may account for how widespread these strains are on plants. Some *Pseudomonas* plant pathogens can also be transmitted through seeds (Praveen Kumar et al. 2015; Abdelfattah et al. 2021), and interestingly, previous work found that pseudomonads were core members of the pea seed microbiome (Chartrel et al. 2021). This suggests that in addition to colonizing plants from the environment, *Pseudomonas* taxa could also be present on germinating pea seeds. Importantly, the prevalence of *Pseudomonadaeceae* within phyllosphere communities correlated with the occurrence of the pyoverdine biosynthesis gene *pvdL*, indicating these taxa are likely capable of producing this fluorescent siderophore. Altogether these results support the conclusion that potentially fluorescent pseudomonads can be abundant as phyllosphere epiphytes and could be influencing aphid behavior.

We found that phyllosphere pseudomonads belong to diverse genetic lineages, with likely implications for function and interactions with aphids. The most commonly identified *Pseudomonas* species in our 16S rRNA gene sequences were *P. fluorescens* (in 14 samples), *P. putida* (13 samples), and *P. syringae* (13 samples). This pattern was also seen in culture isolates, where half belonged to the *P. fluorescens* phylogroup, a similar number belonged with the *P. syringae* phylogroup, and two isolates belonged in the *P. putida* phylogroup. Similarly, the five pseudomonad MAGs recovered also belonged to these three *Pseudomonas* phylogroups. Both *P. putida* and *P. fluorescens* are known to contain plant beneficial taxa (Mercado-Blanco and Bakker 2007; Hol et al. 2013; Berendsen et al. 2015; Costa-Gutierrez et al. 2022). Additionally, we found 12 phyllosphere communities containing *P. rhizophaerae* – a plant beneficial rhizobacterium(Peix et al. 2003; Kwak et al. 2015; Babalola et al. 2021). In contrast, members of the *P. syringae* phylogroup tend to be either commensals or pathogens (Xin et al. 2018; Passera et al. 2019). Previous work found that *P. syringae* strains can be highly infective and virulent to aphids (Smee et al. 2017; Smee et al. 2021), although this may also be true of pseudomonads more broadly (Paliwal et al. 2022). Together, these results support the conclusion that pseudomonads in the *P. sativum* phyllosphere are highly diverse, likely with different functional impacts on plants and aphids.

We found that epiphytic populations of fluorescent pseudomonads elicit a range of behavioral responses from aphids, from avoidance to attraction. About half of the strains tested here caused a significant change in aphid feeding behavior. One of these was the strain *P. syringae* pv. *syringae* B728A, which was previously found to be avoided with 63% of aphid nymphs preferring control leaves to bacterially treated leaves (Hendry et al. 2018). Here we used a different experimental procedure, where whole plants were treated rather than individual leaflets and winged adult aphids were tested. We found higher avoidance with this set up; 73% of aphids preferred control plants to those treated with strain B728A. This difference could be due to fluorescence being more easily detectable when bacteria are present across the whole plant, or due to differences in preferences between nymphs and winged aphids. Alate dispersing aphids show different host preferences than nymphs, and might be more sensitive to color differences (Döring 2014).

A third of the isolates tested for aphid behavior were significantly avoided by aphids, and an additional two isolates showed non significant trends towards avoidance, making some degree of avoidance the most common response (Figure 6). One isolate that showed significant aphid avoidance, #215, is a *P. syringae* isolate like B728a and others used in previous work (Figure 3, Table S3) (Hendry et al. 2018). Potentially this species complex may be more likely to include avoided strains. The other significantly avoided strain, #204, is a *P. rhodesiae* strain (Figure 3, Table S3). This species has been shown to be plant growth promoting and protective against plant pests (Romero et al. 2016; Ye et al. 2022; Mehlferber et al. 2023), suggesting the potential for plant beneficial strains to also protect against aphid pests. These strains had no obvious similarities in the color of their fluorescence in culture (Figure 7) or their siderophore biosynthesis genes (Figure 5) that might be predictive of a tendency to deter aphids.

Surprisingly, an isolate of *P. putida* and an isolate of *P. pergaminensis* were both attractive to aphids (#216 and #220, Figure 6). These isolates were two of the most green-biased fluorescent strains in culture, suggesting that this response could be the result of a sensory bias; aphids are attracted to the color green (Döring and Chittka 2007). Previous work showed that pyoverdine from *P. syringae* pv. *syringae* B728a produces blue fluorescence that aphids avoid (Hendry et al. 2018). The current study also found that strains with higher emissions in the range of wavelengths for the color blue (450 – 500nm) (blue-biased) were avoided by aphids. Together these results set up a potential dichotomy where blue fluorescence is avoided and green fluorescence is attractive, consistent with known aphid responses to these colors (Döring and Chittka 2007; Hendry et al. 2018). However, we do not see a strong correlation between our measures of fluorescence color in culture and aphid behavioral responses. The pattern of attraction was not consistent across all green biased isolates, as a *P. rhodesiae* isolate (#204) was similarly green in culture but caused significant aphid avoidance (Figure 6A, Figure 7). Similarly, we did not see stronger avoidance with more blue biased culture fluorescence, as our most highly avoided isolate *P. syringae* pv. *syringae* strain B728a was not the most blue biased in culture. Isolates of *P. orientalis* and *P. pergaminensis* (#205 and #227) both had more blue biased fluorescence in culture but did not cause significance aphid avoidance. Conditions in media are very different from conditions on plants, which is likely to influence both the type and amount of production for fluorescent compounds like siderophores (Duffy and Défago 1999; Vindeirinho et al. 2021). Therefore, the intensity of fluorescence and color of fluorescence may vary when bacteria are grown on plants compared to growth on iron-limited media.

Sequenced genomes of phyllosphere pseudomonads revealed diversity in the potential to produce various siderophores and other fluorescent compounds. The fluorescent siderophore pyoverdine is considered the primary siderophore produced by most pseudomonads, and our findings confirm this. We found genes for the biosynthesis of pyoverdine in the genomes of all sequenced fluorescent pseudomonas isolates. We also identified orthologs for a pyoverdine binding receptor in most, but not all of these genomes. It is possible, but unlikely that lineages would maintain the ability to produce but not bind pyoverdine (Bruce et al. 2017; Butaitė et al. 2017; Friesen 2020). Instead, this gene may have diverged in some lineages, such as the *P. putida* phylogroup (Figure 5), making it harder to identify with BLAST. In *Pseudomonas aeruginosa*, pyoverdines and their receptors can be under selection for specificity to prevent cheating from non-producing cells (Katiyar and Goel 2004; Smith et al. 2005; Kümmerli et al. 2015; Guadarrama-Orozco et al. 2023), which could drive this divergence. Many isolates (11 out of 23) contained biosynthesis genes for a secondary siderophore as well. Achromobactin, pyochelin, yersiniabactin, and aerobactin, all secondary siderophores known to be produced by pseudomonads, were identified in our isolates. Genes for the production of pyochelin, a fluorescent secondary siderophore known to bind with both iron and zinc ions (Namiranian et al. 1997), were identified in two isolate genomes (Figure 5). Production of this siderophore may enhance or modulate the fluorescence produced by these strains.

Other siderophores identified here are not known to be fluorescent, but the ability to produce other siderophores may influence pyoverdine production rates in nature. Genes for achromobactin biosynthesis were only found in isolates in the *P. syringae* group. Most isolate genomes in this phylogroup possessed both synthesis and receptor genes for achromobactin (Figure 5). Interestingly, orthologs for the achromobactin receptor were common outside of the *P. syringae* group. This receptor is a member of the TonB-dependent receptor family (Berti and Thomas 2009; Ye, Matthijs, et al. 2014), so it is possible that these hits are not specific to achromobactin. Alternatively, achromobactin non-producing lineages in the other phylogroups may be able to cheat and uptake achromobactin produced by other strains (Raaijmakers et al. 1995). We note that strains from the three phylogroups of *P. fluorescens*, *P. putida*, and *P. syringae* often co-occurred in communities, which could lead to selection to uptake non-specific siderophores.

A majority of isolates (14 of 23) possessed a putative gene for the synthesis of the fluorescent pigment phenazine, *phZ*. Phenazine is a secondary metabolite commonly produced by plant-associated pseudomonads (Dar et al. 2020), which can act as a broad spectrum antibiotic (Mavrodi et al. 2013; Perry and Newman 2019) or serve other functions such as improving iron acquisition ability (Wang et al. 2011). Pigmentation of this compound can range from blue, as with pyocyanin, to orange (Dar et al. 2020; Gonçalves and Vasconcelos 2021) and fluorescent emissions range from green to yellow (Zozulya et al. 1997; Wadday et al. 2019), suggesting that its production could influence leaf color similarly to pyoverdine. We did not identify highly similar orthologs for canonical phenazine biosynthesis genes, which are typically found in an operon (Dar et al. 2020), other than *phZ* in our isolates. *Pseudomonas syringae* pv. *tomato* has been reported to produce phenazine, potentially using *phZ* and divergent biosynthesis genes distributed noncanonically across the genome (Li et al. 2016). To our knowledge this has not been confirmed and it is unclear if our isolates could produce this compound.

We found that phyllosphere pseudomonads were capable of producing a range of fluorescent emission colors and intensities. Our survey of emission colors produced by fluorescent phyllosphere isolates revealed a diversity of emission spectra along a range of blue to green biased in color. Pyoverdines, the most common siderophore in our sequenced pseudomonad isolates, produce fluorescence that ranges from blue to yellow (Elliott 1958; Meyer 2000; Tank et al. 2012), as emissions can be altered by structural variability in a peptide chain attached to the conserved chromophore (Budzikiewicz 1997). Therefore, different strains could all be producing pyoverdine but emit different fluorescent spectra. Additionally, differences in fluorescent emissions between strains could be attributed to production of other compounds besides pyoverdine. The emission maximum of unbound pyochelin is 459nm, resulting in a blue fluorescence (Namiranian et al. 1997). Emissions of the two isolates with pyochelin production potential were either blue-biased (#195) or non-biased (#228), although we do not know which siderophores were produced at the time of measurement. As pigmented fluorescent metabolites, phenazines may also influence cumulative fluorescence and visual spectra.

For several isolates tested here aphid behavioral responses showed high variation across replicate experiments. Most isolates had replicate experiments that produced very different outcomes, with differences as high as 33% more aphids than expected on control plants in one replicate and 32% less than expected in another. This wide swing is not due to general variation and noise, because aphids in each replicate did show decisive choices for control versus treated plants. Rather, aphids appear to be making different choices at different times. We speculate that this variation could be caused by bacteria altering the production of fluorescent compounds due to slight changes in environmental conditions on plants. All experiments used plants from the same batch of seeds and the same soil, but it would be difficult to fully control available iron and carbon sources on leaves. Both of these factors can influence whether pyoverdine, or other siderophores, are produced (Bultreys and Gheysen 2000; Husain 2008; Karamanoli et al. 2011; Butaitė et al. 2018; Mendonca et al. 2020; Vindeirinho et al. 2021). Indeed, we find that our isolates varied in color when grown on different carbon sources. Additional environmental factors such as temperature could also be important. For instance, work on a *P. syringae* strain found that pyoverdine production was upregulated at 18°C relative to 28°C (Arvizu-Gómez et al. 2013). Small differences in temperature across replicates could influence the variation that we find.

## Conclusions

Our results suggest that the outcomes of multitrophic interactions between plants, their microbiomes, and aphids are difficult to predict and dependent on environmental context. We find that phyllosphere pseudomonads have the potential to produce a diversity of fluorescent emissions, and so understanding the environmental contexts in which certain compounds are produced is necessary to predict aphid behavioral outcomes. Additionally, this work does not consider volatiles or other compounds potentially produced by bacteria that could be altering aphid behavior. For example, in previous work we demonstrated that avoidance was not a plant mediated effect (Hendry et al. 2018), but we do not control for plant mediated effects in this study. It is possible that colonization by some bacteria could change plant physiology in a way that is attractive to aphids.

Overall, this study shows that fluorescent pseudomonads in the phyllosphere have potential to influence aphid behavior as both attractants and deterrents. Based on these results and previous work we hypothesize that aphid responses to bacterial fluorescence may be driven by underlying sensory biases, as aphids are attracted to green when they are seeking host plants and deterred by blue (Döring and Chittka 2007). This scenario is similar to the idea of an ecological trap, in which biases for certain types of habitat drive animals to choose environments where they ultimately have lower fitness (Hale and Swearer 2016). Such traps have been shown for insects that are attracted to certain colors or polarized light, such as mayflies laying eggs on asphalt roads (Kriska et al. 1998; Horváth et al. 2010; Boda et al. 2014; Acharya et al. 2022). In previous work we found that avoidance of blue fluorescence may be adaptive because it protected aphids from infection by more virulent bacteria (Hendry et al. 2018). With the current work, we show that some isolates can be attractive, but we do not know how this attraction impacts aphid fitness and whether it is an ecological trap. Aphids are susceptible to a diversity of bacterial pathogens, so attraction to bacteria could be bad for aphids (Harada and Ishikawa 1997; Grenier et al. 2006; Stavrinides et al. 2009; Stavrinides et al. 2010; Costechareyre et al. 2012; Paliwal et al. 2022). Even if these biases do not ultimately impact aphid fitness, they can influence aphid distributions and therefore plant health. Understanding these multipartite interactions is therefore important for ecological communities and agricultural ecosystems.

## Acknowledgements

We thank Russell Ligon, Sara Miller, and Michael Sheehan for support with image analysis. Assistance with sampling was provided by Melanie Smee and Catalina Zuluaga Arias, and we thank them, as well as the farm owners and managers Ariana Taylor-Stanley, Lucy Garrison, Chaw Chang, and Betsy Leonard for support in sample collection. Sharon Restrepo, Zahavah Rojer, and Sara Martinez provided much appreciated assistance with DNA extraction. This work was supported by the USDA National Institute of Food and Agriculture, Hatch project no. 1020929, and support to KLH came from the Cornell SUNY Provost Diversity Fellowship.

