## SupplementaryFigures for "Common fluorescent *Pseudomonas* in the phyllosphere can influence aphid behavior in diverse ways"

Supplemental figures

**Figure S1:** Non-metric Multidimensional Scaling (NMDS) based on Bray-Curtis dissimilarities of Family level microbial abundances across phyllosphere metagenomic communities. Points represent metagenomic communities and are colored by the location from which that phyllosphere sample was collected from.

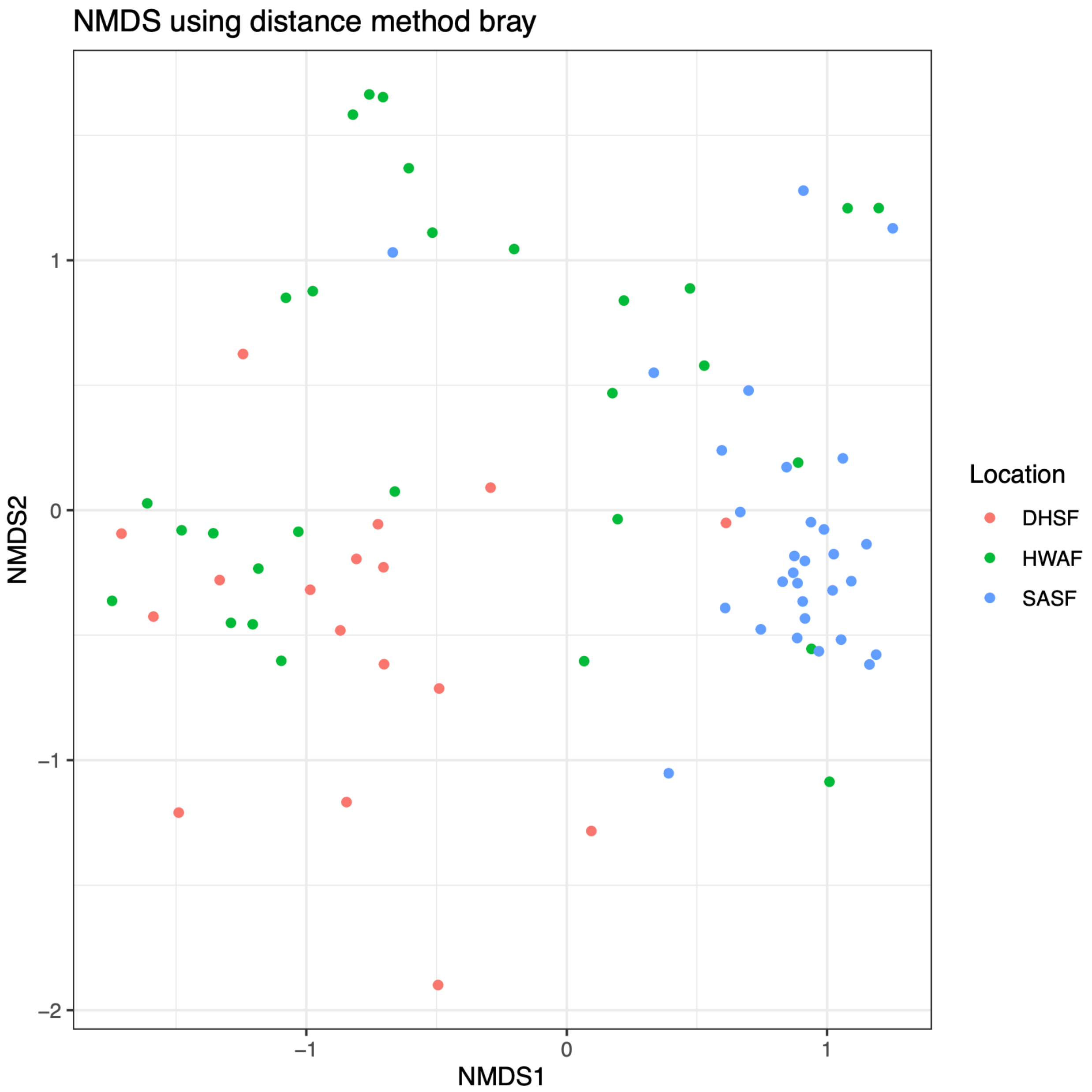

**Figure S2:** Metagenomically Assembled Genomes (MAGs) at the level of Species Genome Bins (SGBs) (right axis) recovered from *Pisum* phyllosphere metagenomes (bottom axis).

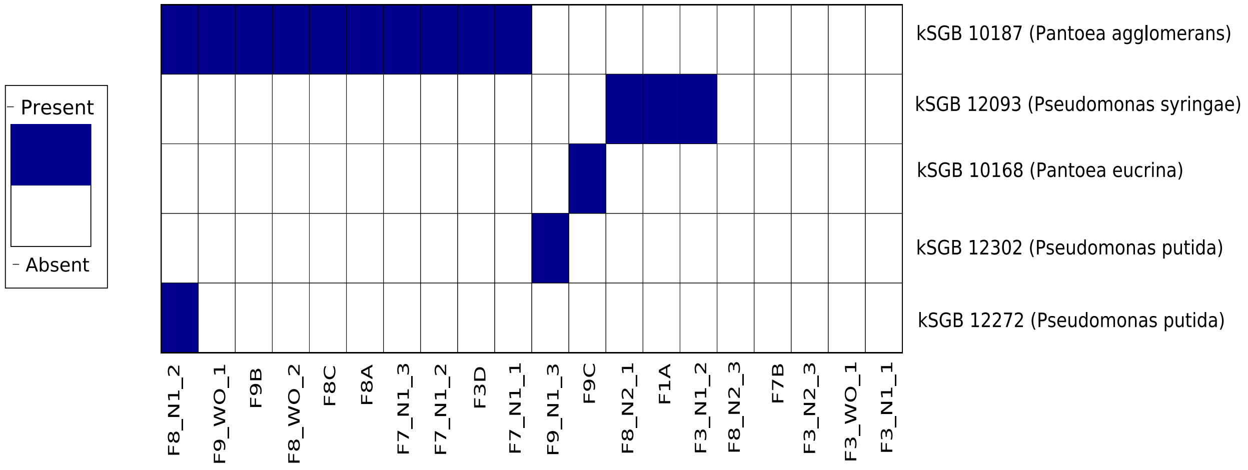

**Supplemental Figure S3** UV reflectance (UVB) and emission intensities in blue and green wavelengths of fluorescent isolates. UVB values were measured alongside blue and green intensities using images of cultured isolates in the same analysis previously described.  Lower UVB values indicate higher UV absorption, which is a proxy for higher fluorescence intensity.

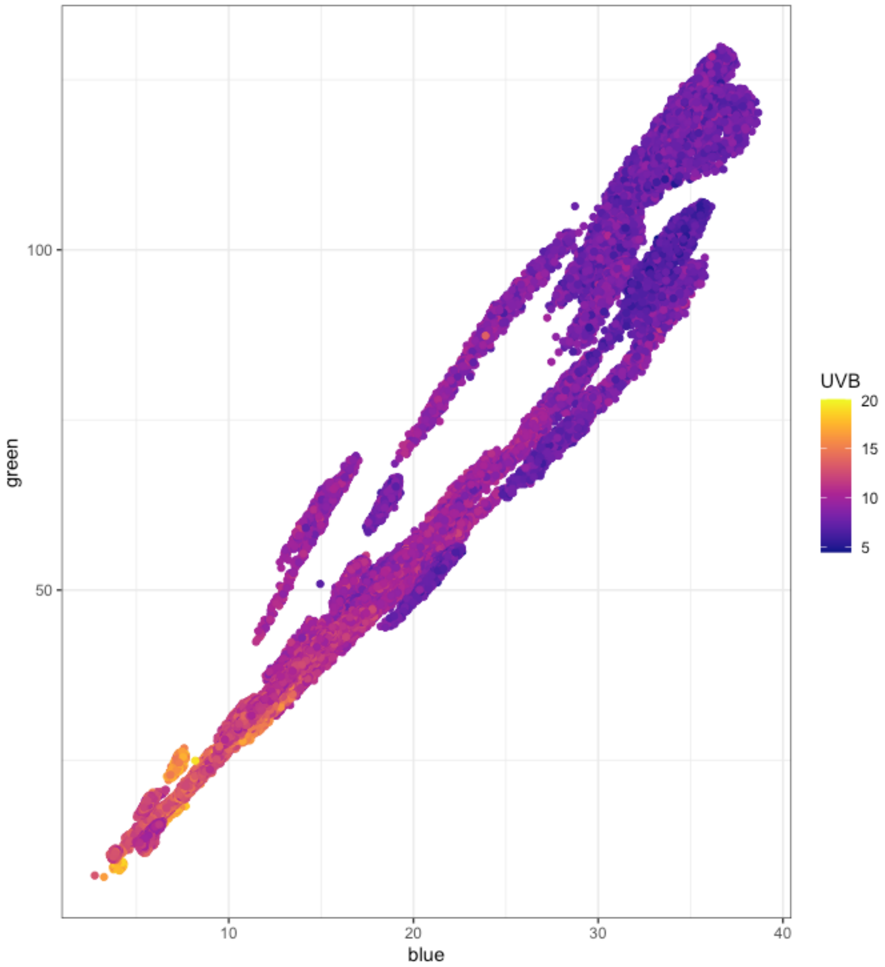

**Supplemental Figure S4. Fluorescence in different conditions. A)** Normalized ratios of emission intensities in blue (380nm excitation, 430nm emission) and green (400nm excitation, 530 nm emission) wavelengths emitted by isolate #194 grown in *Pseudomonas* Minimal Media (PMM) with either 0.4% glucose (bottom bar) or 1.5% glycerol (right bar for each strain) as the sole carbon source. Emission values of strains over baseline emissions of media alone were normalized to cell growth using the optical density of each strain ($OD_{600}$). Values shown are the proportions of normalized blue and green emissions out of the total emission. **B)** Qualitative image of diverse fluorescence spectra of isolate #194 when grown overnight in liquid PMM supplemented with distinct carbon sources. From left to right, cultures are grown PMM without glycerol (left tube), with 0.4% glucose, with 0.4% sucrose, with 0.4% fructose, or with 0.4% glycerol (right tube).

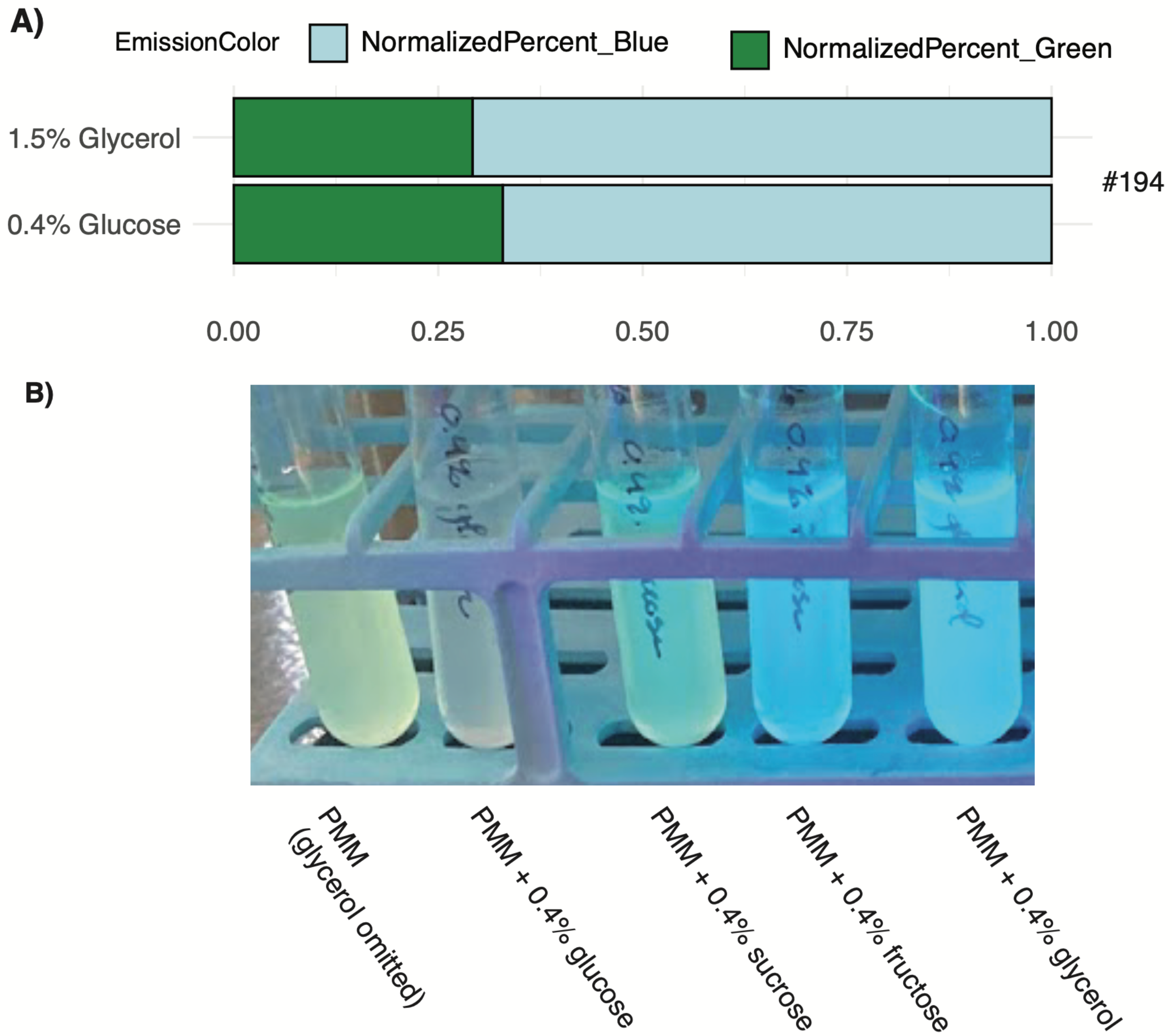

**Supplemental Table S1**: Reads/coverage/assembly stats for samples/MAGs/genomes accession numbers.

**Supplementary table S2:** Siderophore and phenazine gene queries used to confirm fluorescent isolate annotations via NBCI BLAST.

**Supplemental Table S3:** Best match of fluorescent phyllosphere isolates and their nearest relatives by average nucleotide similarity based on BLAST+ (percent ANIb).

**Supplemental table S4:** Results of generalized linear mixed-effects models (GLMMs) used to evaluate the significance of aphid choice. Date and experimental replicate were included as random effects, with strain identity as the fixed effect. Likelihood ratio tests were used to determine whether strain identity significantly affected aphid choice compared to the null expectation. The model intercept was removed to test for deviations from our null expectation by including -1 as a variable. The resulting p value indicates if the probability of aphids choosing control plants significantly varied from 50%.

| *Strain* | *Estimate* | *Std. Error* | *z value* | *Pr(>\|z\|)* | *Significance* |
|---|---|---|---|---|---|
| *#194* | 0.2612 | 0.28578 | 0.914 | 0.360727 | |
| *#204* | 0.85678 | 0.30655 | 2.795 | 0.005192 | ** |
| *#205* | 0.23072 | 0.3366 | 0.685 | 0.493058 | |
| *#215* | 0.55246 | 0.28101 | 1.966 | 0.049299 | * |
| *#216* | -0.63111 | 0.29503 | -2.139 | 0.032422 | * |
| *#220* | -1.32875 | 0.35486 | -3.744 | 0.000181 | *** |
| *#227* | 0.37784 | 0.26755 | 1.412 | 0.157875 | |
| *#228* | 0.03401 | 0.27831 | 0.122 | 0.902733 | |
| *B728A* | 1.0666 | 0.43773 | 2.437 | 0.014823 | * |

*Signif. codes: 0 '***' 0.001 '**' 0.01 '*' 0.05 '.' 0.1 ' ' 1*

**Supplemental table S5:** Results of pairwise Permutational Multivariate Analysis of Variances (PERMANOVA) testing for variations in phyllosphere microbial communities between sampling locations. Test was performed using Bray Curtis dissimilarities and individual host plant was included as a nested variable. Location codes are as follows; HWAF= Here We Are Farm, SASF = Stick And Stone Farm, DHSF = Dilmun Hill Student Farm.

| | *location 1* | *location 2* | *p* | *p.adj* |
|---|---|---|---|---|
| *comparison 1* | HWAF | DHSF | 0.02 | 0.03 |
| *comparison 2* | HWAF | SASF | 0.01 | 0.03 |
| *comparison 3* | DHSF | SASF | 0.03 | 0.03 |
